# Non-linear transfer characteristics of stimulation and recording hardware account for spurious low-frequency artifacts during amplitude modulated transcranial alternating current stimulation (AM-tACS)

**DOI:** 10.1101/246710

**Authors:** Florian H. Kasten, Ehsan Negahbani, Flavio Fröhlich, Christoph S. Herrmann

**Affiliations:** Experimental Psychology Lab, Department of Psychology, European Medical School, Cluster for Excellence “Hearing for All”, Carl von Ossietzky University, Oldenburg, Germany; Neuroimaging Unit, European Medical School, Carl von Ossietzky University, Oldenburg, Germany; Department of Psychiatry, University of North Carolina, Chapel Hill, NC, USA; Department of Cell Biology and Physiology, University of North Carolina, Chapel Hill, NC, USA; Department of Biomedical Engineering, University of North Carolina, Chapel Hill, NC, USA; Neuroscience Center, University of North Carolina, Chapel Hill, NC, USA; Department of Neurology, University of North Carolina, Chapel Hill, NC, USA; Carolina Center for Neurostimulation, University of North Carolina, Chapel Hill, NC, USA; Research Center Neurosensory Science, Carl von Ossietzky University, Oldenburg, Germany

**Keywords:** amplitude modulated transcranial alternating current stimulation (AM-tACS), MEG, EEG, artifact, tACS, stimulation hardware

## Abstract

Amplitude modulated transcranial alternating current stimulation (AM-tACS) has been recently proposed as a possible solution to overcome the pronounced stimulation artifact encountered when recording brain activity during tACS. In theory, AM-tACS does not entail power at its modulating frequency, thus avoiding the problem of spectral overlap between brain signal of interest and stimulation artifact. However, the current study demonstrates how weak non-linear transfer characteristics inherent in stimulation and recording hardware can reintroduce spurious artifacts at the modulation frequency. The input-output transfer functions (TFs) of different stimulation setups were measured. The setups included basic recordings of signal-generator and stimulator outputs as well as M/EEG phantom measurements. 6^th^-degree polynomial regression models were fitted to model the input-output TFs of each setup. The resulting TF models were applied to digitally generated AM-tACS signals to predict the location of spurious artifacts in the spectrum. All four setups measured for the study exhibited low-frequency artifacts at the modulation frequency and its harmonics when recording AM-tACS. Fitted TF models showed non-linear contributions significantly different from zero (all p < .05) and successfully predicted the frequency of artifacts observed in AM-signal recordings. Results suggest that even weak non-linearities of stimulation and recording hardware can lead to spurious artifacts at the modulation frequency and its harmonics. Thus, findings emphasize the need for more linear stimulation devices for AM-tACS and careful analysis procedures, which take into account these low-frequency artifacts to avoid confusion with effects of AM-tACS on the brain.

## 1 Introduction

Transcranial alternating current stimulation (tACS) is receiving growing popularity as a tool to interfere with endogenous brain oscillations in a frequency specific manner (Fröhlich and McCormick’ 2010; Helfrich et al., 2014; Herrmann et al., 2013; Ozen et al., 2010; Zaehle et al., 2010), allowing to study causal relationships between these oscillations and cognitive functions (Fröhlich, 2015; Herrmann et al., 2016). Further, its use might offer promising new pathways for therapeutic applications to treat neurological or psychiatric disorders associated with dysfunctional neuronal oscillations (Brittain et al., 2013; Herrmann and Demiralp, 2005; Uhlhaas and Singer, 2012, 2006).

While mechanisms of tACS have been studied in animals (Fröhlich and McCormick, 2010; Kar et al., 2017; Ozen et al., 2010; Reato et al., 2010) and using computational modelling (Ali et al., 2013; Reato et al., 2010; Zaehle et al., 2010), the investigation of tACS effects in human subjects has so far mostly been studied behaviorally (Kar and Krekelberg, 2014; Lustenberger et al., 2015; Neuling et al., 2012), by measuring BOLD response (Cabral-Calderin et al., 2016; Violante et al., 2017; Vosskuhl et al., 2016), or by tracking outlasting effects in M/EEG signals (Kasten et al., 2016; Kasten and Herrmann, 2017; Neuling et al., 2013; Veniero et al., 2015; Vossen et al., 2015; Zaehle et al., 2010). Due to a strong electro-magnetic artifact, which spectrally overlaps with the brain oscillation under investigation, online measurements of tACS effects in M/EEG is challenging. However, uncovering these online effects is crucial as the aforementioned approaches can only provide limited, indirect insights to the mechanisms of action during tACS in humans. In addition, online monitoring of physiological signals during stimulation may enable closed-loop applications that can provide potentially more powerful, individually tailored, adaptive stimulation protocols (Bergmann et al., 2016). Some authors applied artifact suppression techniques such as template subtraction (Dowsett and Herrmann, 2016; Helfrich et al., 2014; Voss et al., 2014) or spatial filtering (Neuling et al., 2015; Ruhnau et al., 2016) to recover brain signals obtained during concurrent tACS-M/EEG. However, these approaches are computationally costly, and therefore i.e. difficult to implement in closed-loop protocols. Further, their application is limited as they fail to completely suppress the artifact and analysis approaches must be limited to robust procedures to avoid false conclusions about stimulation effects (Neuling et al., 2017; Noury et al., 2016; Noury and Siegel, 2017).

As a solution to these issues, amplitude modulated tACS (AM-tACS), using a high frequency carrier signal which is modulated in amplitude by a lower frequency modulation signal, chosen to match the targeted brain oscillation has been proposed (Witkowski et al., 2016). Amplitude modulated signals contain spectral power at the frequency of the carrier signal (*f_c_*; and two sidebands at *f_c_* ± *f_m_;* modulation frequency), but no power at *f_m_* itself (see Figure 1 for an illustration). Consequently, the tACS artifact would be shifted into higher frequencies, elegantly avoiding spectral overlap with the targeted brain oscillation. However, more recently low-frequency artifacts at *f_m_* have been reported in sensor-level MEG recordings during AM-tACS (Minami and Amano, 2017). These artifacts required the application of advanced artifact suppression algorithms (Minami and Amano, 2017). Although the authors of that study explained these artifacts by non-linear characteristics of the digital-analog conversion, a detailed investigation into these low-frequency artifacts arising during AM-tACS and how these emerge has not yet been provided. In fact, the process of stimulation on the one side and signal recording on the other side involves at least one step of digital-analog (generating a stimulation signal) and one step of analog-digital conversion (sampling brain signal plus stimulation artifact). The linearity of these conversions, however, is naturally limited by properties of the hardware in use (Vargha et al., 2001). To further complicate the situation, the amplification involved in the recording process using M/EEG can be another potential source of nonlinearity. The amplitudes usually applied in tACS can potentially cause signals/artifacts, beyond the dynamic range where the measurement devices exhibit linear transfer characteristics (Cooper, R., Osselton, J. W., & Shaw, 1974). In general all electronic components, including those that are usually idealized as being linear (i.e. resistors), exhibit some degree of non-linearity in reality, especially when operating under extreme conditions (Maas, Stephen, 2003).

To shed more light on the effects of non-linearity of stimulation and recording hardware on AM-tACS signals, input-output transfer functions (TFs) of different AM-tACS setups were estimated and evaluated with respect to their performance in predicting low-frequency artifacts of AM-tACS^1^.

## 2 Materials & Methods

In order to characterize non-linearities inherent in different tACS setups, the transfer functions (TFs) relating input-output amplitudes of four different tACS setups, with increasing complexity, were recorded and modeled by polynomial regression models. Additionally, AM-tACS signals were recorded to demonstrate the presence of low-frequency artifacts. TF models were applied to digital AM-signals to predict output spectra of the physical recordings. The following four setups were evaluated. No human or animal subjects were involved in the experiment.

### 2.1 Test Setups

#### 2.1.1 Basic DAC recording

For the first, basic setup, a digital/analog-analog/digital converter (DAC; NiUSB-6251, National Instruments, Austin, TX, USA) recorded its own output signal. The signal was digitally generated using Matlab 2016a (The MathWorks Inc., Natick, MA, USA) and streamed to the DAC via the Data Acquisition Toolbox. The signal was generated and recorded at a rate of 10 kHz (Figure 1A).

**Figure 1.**
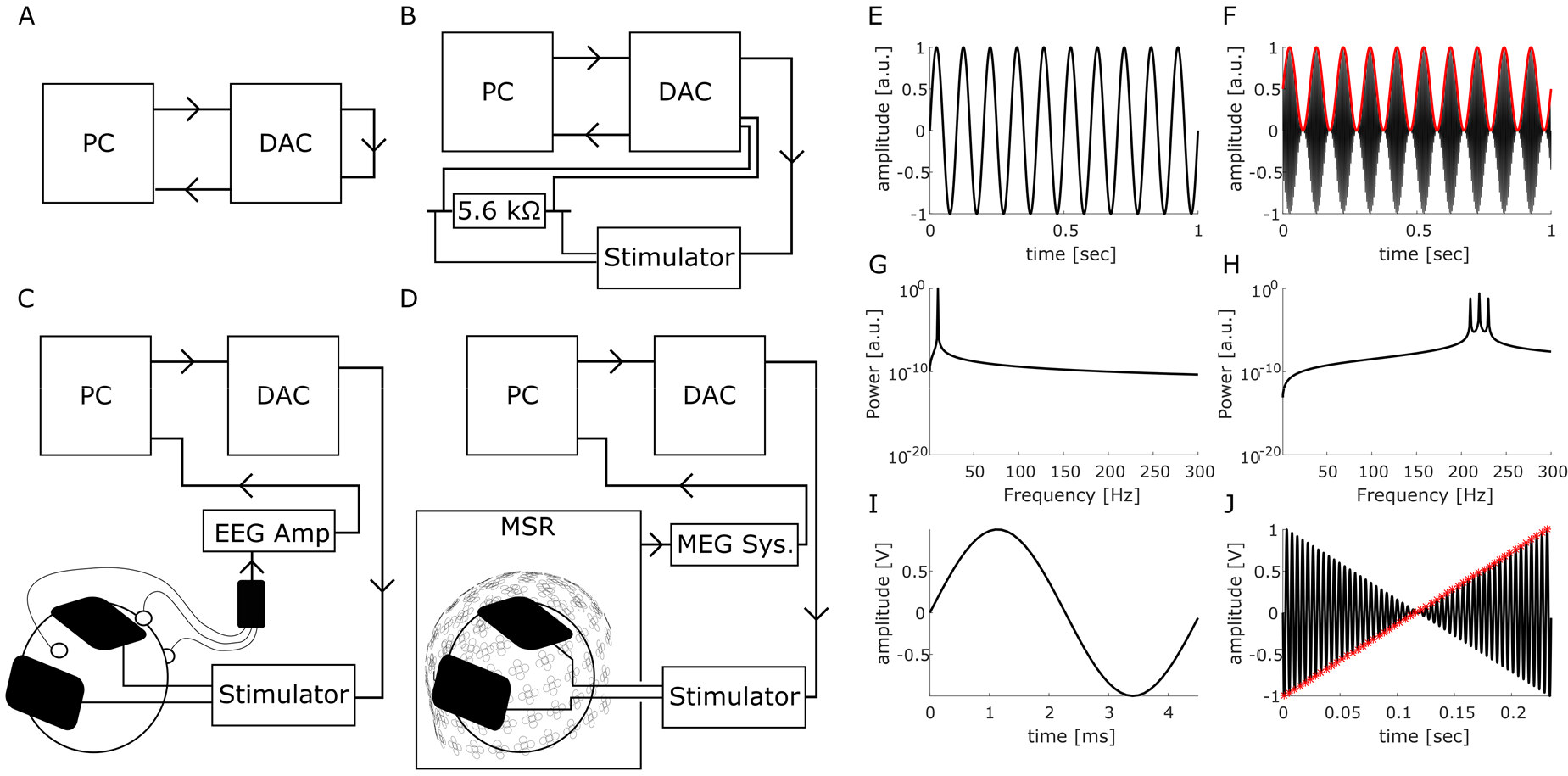
Experimental setups and signals. **(A-D)** Schematic representations of the evaluated setups. For details refer to the “**Test setups**” section in the manuscript. DAC: Digital-Analog converter. MSR: Magnetically shielded room. Arrows indicate the direction of signal flow (**E,F**) Time domain representations of a low-frequency sine-wave classically used for tACS (**E**) and an amplitude modulated sine wave with a carrier frequency of 220 Hz modulated at 10 Hz (**F**). Red curve depicts the 10 Hz envelope of the signal. (**G,H**) Frequency-domain representations of the tACS signals. While the 10 Hz sine wave exhibits its power at 10 Hz (**G**), the amplitude modulated signal only exhibits power at the carrier frequency and two side-bands, but no power at the modulation frequency (**F**). (**I**) Probe stimulus for measuring the setups transfer curves was a 220 Hz single-cycle sine wave. Probe stimuli of different amplitude were concatenated to a sweep (**J**). Red asterisks mark points that were extracted as *V_out_* measure. To enhance visibility of the general concept, a sweep consisting of 51 probes is displayed here. For the actual measurements of the TFs 10 sweeps with 10001 probes were used.

#### 2.1.2 DAC & tACS stimulator

In the second setup the DAC was connected to the remote input of a battery-driven constant current stimulator (DC Stimulator Plus, Neuroconn, Illmenau, Germany). Stimulation was administered to a 5.6 kQ resistor. The signal was recorded from both ends of the resistor using the DAC (Figure 1B).

#### 2.1.3 DAC & tACS recorded from phantom using EEG

In the third setup the DC Stimulator was connected to two surface conductive rubber electrodes attached to a melon serving as a phantom head. Electrodes were attached using an electrically conductive, adhesive paste (ten20, Weaver & Co., Aurora, CO, USA). The signal was recorded from an active Ag/AgCl EEG electrode (ActiCap, Brain Products, Gilching, Germany), placed between the tACS electrodes. Two additional electrodes were attached to the phantom to serve as reference and ground for the recording (positions were chosen to mimic a nose-reference and a ground placed on the forehead). The signal was generated by the DAC at a rate of 10 kHz and recorded at 10 kHz using a 24-bit ActiChamp amplifier (Brain Products, Gilching, Germany). EEG and stimulation electrode impedances were kept below 10 kQ (Figure 1C).

#### 2.1.4 DAC & tACS recorded from phantom using MEG

Finally, the phantom was recorded using a 306-channel whole-head MEG system (Elekta Neuromag Triux, Elekta Oy, Helsinki, Finland) located inside a magnetically shielded room (MSR; Vacuumschmelze, Hanau, Germany). Signals were recorded without internal active shielding at a rate of 1 kHz and online filtered between 0.3 and 330 Hz. The stimulation signal was gated into the MSR via the MRI-extension kit of the DC stimulator (Neuroconn, Illmenau, Germany; Figure 1D).

### 2.2 Transfer function and AM-tACS measurements

A probe stimulus consisting of a one cycle sine wave at 220 Hz was used to obtain measurements of each setups transfer function (TF). 10001 probes of linearly spaced amplitudes *(V¿„)*, ranging from −10 V to 10 V for the first setup, from −0.75 V to 0.75 V for the second and third setup, and from −0.5 V to 0.5 V for the MEG setup, were concatenated to a sweep stimulus with a total duration of approximately 45 sec. (see Figure 1I-J for a schematic visualization). Amplitudes had to be adjusted for setups involving the DC stimulator to account for higher output voltages due to the 2 mA per V voltage-to-current conversion of the remote input. The chosen input voltages correspond to a maximum output of 3 mA peak-to-peak amplitude of the DC stimulator (a maximum current of 2 mA was chosen for the MEG setup to avoid saturation and flux trapping of MEG sensors). Ten consecutive sweeps were applied and recorded for each setup. In order to evaluate how well the obtained TF can predict artifacts in the spectrum of AM-tACS, AM-signals with *f_c_* = 220 Hz and *f_m_* = 10 Hz, 11 Hz, and 23 Hz at different amplitudes (100%, 66.7%, 33.4% and 16.16% of the maximum range applied during the TF recording) were generated. Amplitudes were chosen to produce output currents of 3 mA, 2 mA, 1 mA, and 0.5 mA when using the DC-Stimulator (2 mA, 1.3 mA, 0.66 mA, 0.33 mA for the MEG setup). AM-signals were computed based on the following equation:

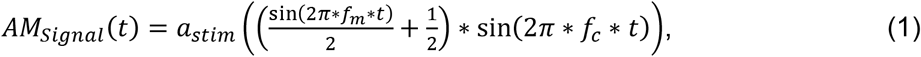

where *a_stim_* is the stimulation amplitude, *f_m_* is the modulation frequency and *f_c_* is the carrier frequency. Each signal was generated and recorded with 60 repetitions to increase signal-to-noise ratios.

### 2.3 Data Analysis

Data analysis was performed using Matlab 2016a (The MathWorks Inc., Natick, MA, USA). The fieldtrip toolbox (Oostenveld et al., 2011) was used to import and segment M/EEG recordings. All scripts and underling datasets are available online (https://osf.io/czb3d/).

#### 2.3.1 Data processing and transfer function estimation

The recorded sweeps were epoched into segments containing single cycles of the sine-waves used as probes. All Segments were baseline corrected and the peak-amplitude *(V_out_)* of each epoch was extracted by identifying the minimum (for *V_in_* < 0) or maximum values (for *V_in_ ≥ 0)* within each segment. A 6^th^-degree polynomial regression model was fitted to each repetition of the sweep to predict *V_out_* (recorded peak amplitudes) as a function of *V_in_* (generated peak amplitudes) using a least-square approach:

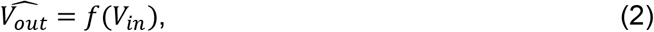

with:

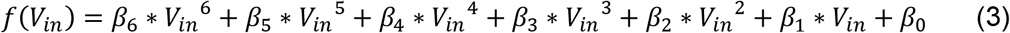

The fitting procedure was performed separately for each sweep to obtain measures of variance for each of the coefficients. Coefficients were averaged subsequently and the resulting function was used to model each systems TF. R^2^-values were calculated as measures for goodness of fit.

In order to evaluate the performance of the TF models in predicting low-frequency AM-tACS artifacts of the setups, the digitally generated AM-tACS signals were fed through the TF models. Subsequently, the predicted output signals were compared to the AM-tACS recordings acquired for each setup. To this end, power spectra of the original digital, the predicted and the recorded AM-signals were computed. The resulting power spectra of the AM-signals were averaged over the 60 repetitions. For the MEG recording, results are presented for an exemplary parieto-occipital gradiometer sensor (MEG2113).

#### 2.3.2 Identification of low-frequency artifacts

To identify systematic artifacts in the spectrum of the AM-signal in the noisy recordings, the averaged power spectra were scanned for artifacts within a range from 2 Hz to 301 Hz. Artifacts were defined as the power at a given frequency being altered by at least 5% as compared to the mean power of the two neighboring frequencies. The identified artifacts were statistically compared to the power in the two neighboring frequencies using student’s t-tests. Bonferroni-correction was applied to strictly account for multiple comparisons.

#### 2.3.3 Simulation

To evaluate the effect of each non-linear term in the TF models on the output signal, a simulation was carried out. To this end an amplitude modulated signal with *f_c_* = 220 Hz and *f_m_* = 10 *Hz* was evaluated by simplified TFs where all coefficients were set to zero except for the linear and one additional non-linear term which were set to one in each run. This procedure leads to exaggerated output spectra that do not realistically resemble the recorded TFs. However, they are well suited to illustrate the spectral artifacts arising from each of the non-linear terms.

In addition to the AM-signal, we generated a temporal interference (TI) signal that was recently proposed as a tool to non-invasively stimulate deep structures of the brain (Grossman et al., 2017). TI stimulation consists of two externally applied, high frequency sine waves of slightly differing frequencies that result in an AM-signal where their electric fields overlap. Since the generation of this AM-signal is mathematically slightly different as compared to the other AM-tACS approach, this signal was separately modelled for two stimulation signals based on the following equation:

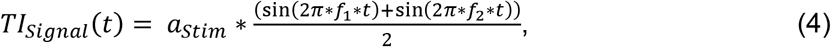

with *f = 200 Hz* and *f*_2_ = 210 *Hz.* The overlap of these two frequencies results in an amplitude modulation at 10 Hz.

## 3 Results

### 3.1 Systematic artifacts at modulation frequency of AM-tACS and harmonics

Analysis of the AM-tACS recordings identified systematic artifacts at the modulation frequency and its harmonics that statistically differed from power at neighboring frequencies in all setups (all p < .05; Figure 2 and 3). Notably, these artifacts were comparatively small, albeit still significant at larger amplitudes, when the DAC measured its own output without any further devices in the setup (Figure 2 **left).** When the complexity of the setup was increased, more and stronger artifacts were observed (Figure 2 right, Figure 3). The number and size of artifacts also tended to increase with stronger stimulation amplitudes. Figures 2 and 3 depict lower frequency spectra (1 Hz - 50 Hz) for all setups and frequency-amplitude combinations tested.

### 3.2 Setups exhibit non-linear transfer characteristics

To obtain a model of each setups TF, 6^th^-degree polynomial regression models were fitted to the input-output amplitudes of the probe stimuli. All setups tested in this study exhibited coefficients of the non-linear terms of the fitted TFs significantly differing from zero. In setups 1, 2, and 4 all model coefficients significantly differed from zero (all *p* < .004; bonferroni corrected). For the EEG setup, coefficients *ß*_2_ (p < .02), *ß*_5_ (p < .004) and *ß*_6_ (p < .007) significantly differed from zero. Results are summarized in **Table 1.** High goodness of fit values were achieved for all setups under investigation (*R^2^* > .99), indicating that the polynomial functions provide powerful models to describe the input-output characteristics of the setups. Importantly, the non-linearities found during this analysis are subtle compared to the contribution of the linear terms in each TF. This leads to the impression of linearity when visually inspecting each setups TF (Figure 2, 3 **top panel).** However, as it will be shown in the following, these small deviations from linearity are sufficient to cause the low frequency artifacts observed during the AM-tACS recordings.

**Figure 2.**
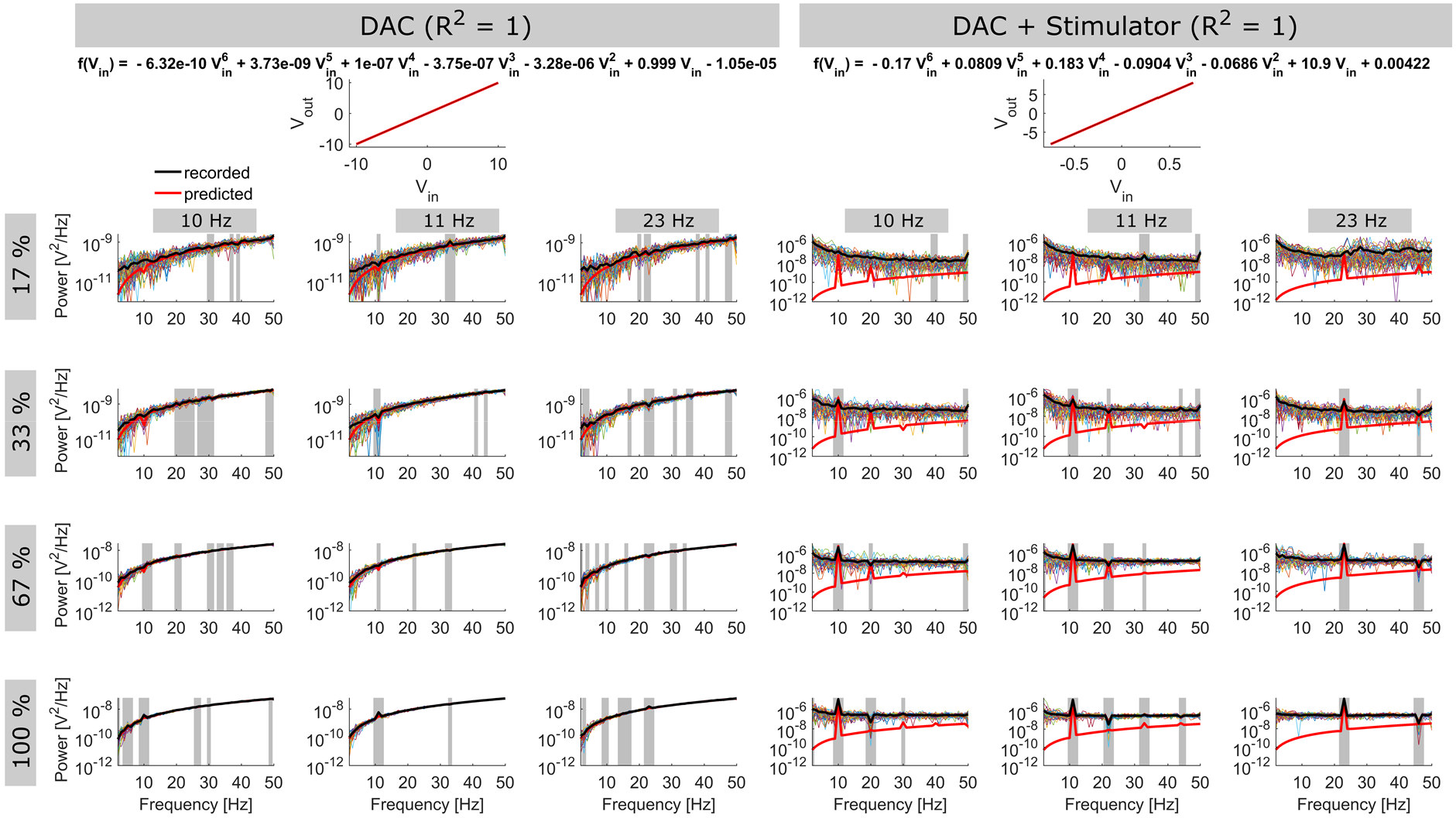
Transfer functions (top row) and spectra (lower rows) of setups of the DAC and Stimulator setup. TFs **(top)** show recorded probe stimulus amplitudes in relation to their input amplitudes (*V_out_/V_in_*; black dots), as well as the course of the TF model (red line). The corresponding function is displayed in the title. Spectra show average power at each frequency in the different AM-recordings (black line). Thin colored lines show power spectra for each of the 60 repetitions. Red line shows the spectrum predicted by evaluating the digital AM-signal by the estimated TF of the setup. Grey areas indicate frequencies significantly differing in power compared to the two neighboring frequencies (p < .05, bonferroni corrected). Please note the different scaling of the power spectra. To enhance visibility, spectra are limited to the frequency range between 1 Hz and 50 Hz. Please refer to the **Supplementary Materials** for an alternative version of the figure, covering the full frequency range between 1 and 300 Hz.

**Table 1.** Transfer function coefficients tested for deviation from zero. Coefficients of the 10 polynomial functions fitted for each setups TF recordings were tested against zero using student’s t-test (two-sided, bonferroni corrected). Mean and standard deviation are shown for each coefficient.

|  | *Mean* | *Std.* | *df* | *T* | *p* |
| --- | --- | --- | --- | --- | --- |
| **DAC** |  |  |  |  |  |
| $\beta_0$ | -1.05e-05 | 4.80e-06 | 9 | -6.92 | < .001* |
| $\beta_1$ | 0.9988 | 1.86e-05 | 9 | > 100 | < .001* |
| $\beta_2$ | -3.28e-06 | 7.02e-07 | 9 | -14.79 | < .001* |
| $\beta_3$ | -3.75e-07 | 7.16e-08 | 9 | -16.56 | < .001* |
| $\beta_4$ | 9.99e-08 | 2.31e-08 | 9 | 13.69 | < .001* |
| $\beta_5$ | 3.73e-09 | 5.77e-10 | 9 | 20.41 | < .001* |
| $\beta_6$ | -6.32e-10 | 1.72e-10 | 9 | -11.63 | < .001* |
| **DAC + Stimulator** |  |  |  |  |  |
| $\beta_0$ | 0.0042 | 0.0009 | 9 | 15.37 | < .001* |
| $\beta_1$ | 10.8640 | 0.0123 | 9 | > 100 | < .001* |
| $\beta_2$ | -0.0686 | 0.0153 | 9 | -14.14 | < .001* |
| $\beta_3$ | -0.0904 | 0.0324 | 9 | -8.83 | < .001* |
| $\beta_4$ | 0.1838 | 0.0606 | 9 | 9.54 | < .001* |
| $\beta_5$ | 0.0809 | 0.0484 | 9 | 5.28 | < .001* |
| $\beta_6$ | -0.1702 | 0.0712 | 9 | -7.56 | < .001* |
| **EEG** |  |  |  |  |  |
| $\beta_0$ | -0.0001 | 0.0001 | 9 | -5.27 | < .001* |
| $\beta_1$ | 0.1736 | 0.0007 | 9 | > 100 | < .001* |
| $\beta_2$ | 0.0024 | 0.0017 | 9 | 4.44 | .002* |
| $\beta_3$ | -0.0006 | 0.0024 | 9 | -0.81 | .44 |
| $\beta_4$ | 0.0035 | 0.0069 | 9 | 1.64 | .14 |
| $\beta_5$ | -0.0058 | 0.0035 | 9 | -5.30 | < .001* |
| $\beta_6$ | -0.0118 | 0.0078 | 9 | -4.80 | .001* |
| **MEG** |  |  |  |  |  |
| $\beta_0$ | -0.0009 | 0.0002 | 9 | -16.35 | < .001* |
| $\beta_1$ | 11.3235 | 0.0576 | 9 | > 100 | < .001* |
| $\beta_2$ | 0.0267 | 0.0121 | 9 | 6.97 | < .001* |
| $\beta_3$ | 0.3033 | 0.0393 | 9 | 24.41 | < .001* |
| $\beta_4$ | -0.5931 | 0.1532 | 9 | -12.24 | < .001* |
| $\beta_5$ | -1.1228 | 0.2065 | 9 | -17.19 | < .001* |
| $\beta_6$ | 2.1034 | 0.5192 | 9 | 12.81 | < .001* |

### 3.3 Transfer functions predict frequency of spurious artifacts

When applying the TF models to the digital AM-signals, the resulting spectra provide accurate predictions of the systematic low-frequency artifacts at *f_m_* of the AM-signal and its lower harmonics in the recordings. For the first two setups, where the TF models’ goodness of fit is equal to 1, the predicted spectra also capture the amplitudes low-frequency artifacts with relatively high accuracy (Figure 2). For the two later setups, however, the predicted spectrum apparently underestimates amplitudes of the artifacts (Figure 3). In summary, results suggest that the polynomial functions fitted to the data successfully captured the non-linear process leading to the low-frequency artifacts at *f_m_*, although for the later setups, that exhibited more noise during the measurements, accuracy of the fits seems not sufficient to accurately predict the artifacts amplitudes. In addition, it should be noted that the application of a TF to a pure digital AM-signal can never completely capture the effects of the recording process that involves measurement of noise and external interferences (i.e. line-noise).

**Figure 3.**
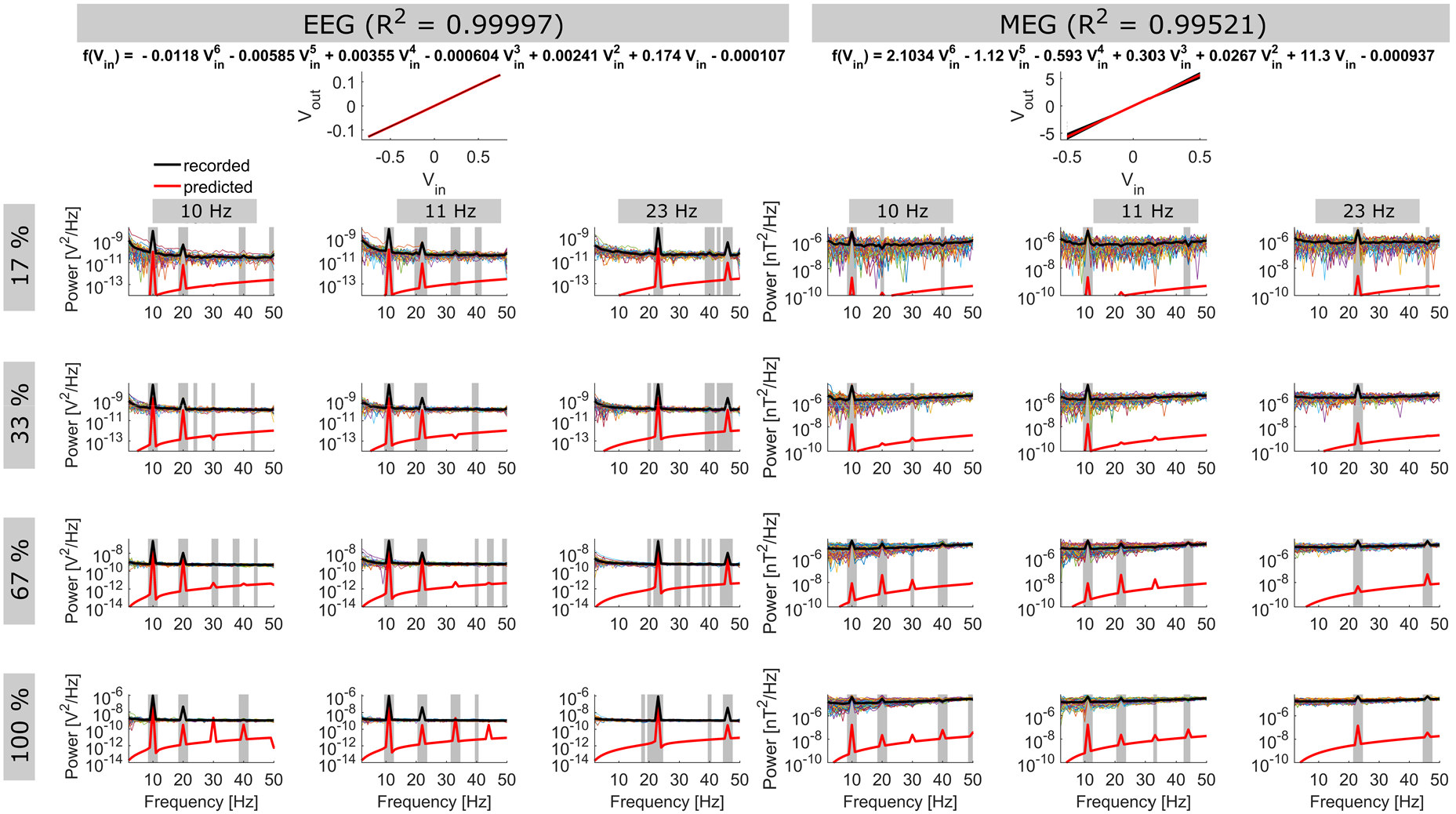
Transfer functions (top row) and spectra (lower rows) of the EEG and MEG setup. TFs **(top)** show recorded probe stimulus amplitudes in relation to their input amplitudes (V_out_/V_in_; black dots), as well as the course of the TF model (red line). The corresponding function is displayed in the title. Spectra show average power at each frequency in the different AM-recordings (black line). Thin colored lines show power spectra for each of the 60 repetitions. Red line shows the spectrum predicted by evaluating the digital AM-signal by the estimated TF of the setup. Grey areas indicate frequencies significantly differing in power compared to the two neighboring frequencies (p < .05, bonferroni corrected). Please note the different scaling of the power spectra. To enhance visibility, spectra are limited to the frequency range between 1 Hz and 50 Hz. Please refer to the **Supplementary Materials** for an alternative version of the figure, covering the full frequency range between 1 and 300 Hz.

### 3.4 Simulating the isolated effect of non-linear TF-terms

Based on the results presented so far, it was possible to characterize each the non-linearity of each setup and to demonstrate that the estimated TF can be used to predict artifacts in the recorded AM-signals. However, since the obtained TFs are rather complex, a simulation was carried out to investigate the artifacts caused by each of the non-linear terms in isolation. The spectra obtained from this simulation are depicted in Figure 4. While a solely linear TF does not change the spectral content of the AM-signal at all (Figure 4 **top left)** polynomial terms with odd exponents > 1 result in additional side bands around *f_c_* of the AM-signal (Figure 4 **middle, bottom left).** In contrast, terms with even exponents induced artifacts at *f_m_* and its harmonics (Figure 4 **right column).** The higher the exponent of the polynomial terms the more sidebands and higher harmonics are introduced to the spectrum, respectively. A separate simulation for an AM-signal resulting from temporal inteference (Grossman et al., 2017) yielded a similar result (Supplementary Figure S3).

**Figure 4.**
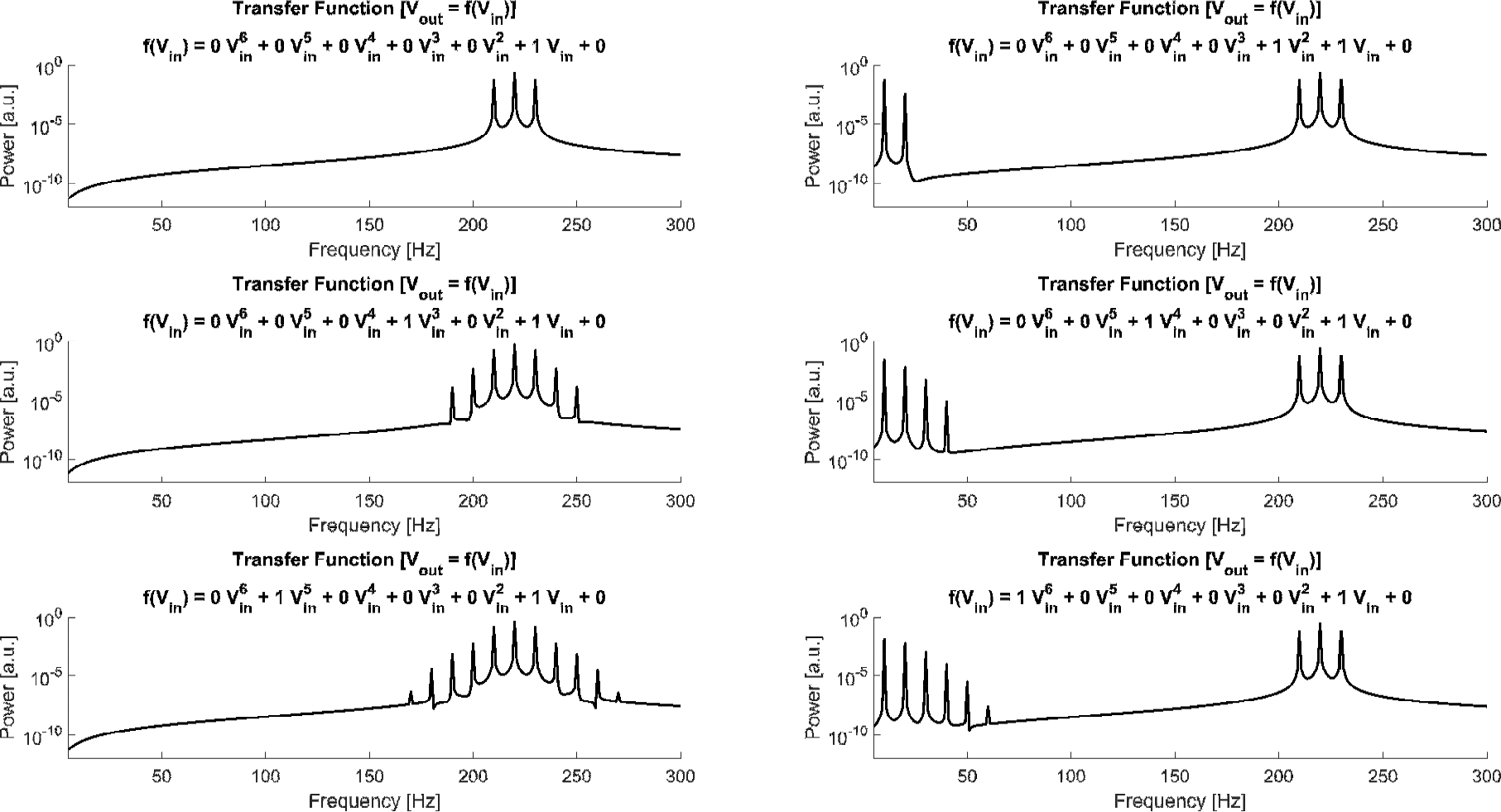
Simulation results. (Left column) Spectra resulting from evaluating the digital AM-signal using a simplified TF. A solely linear TF (**top left**) perfectly resembles the input spectrum. Setting the coefficient of an additional polynomial term with an odd-valued exponent to 1 resulted in additional side bands around fc (**middle** and **bottom left**). In contrast, setting the coefficient of an additional polynomial term with an even-valued exponent to 1 resulted in artifacts at *f_m_* and its harmonics (**right column**). The higher the exponent of the polynomial terms, the more side bands/harmonic artifacts they introduced. The polynomial function applied to generate each spectrum is printed on top of each plot.

## 4 Discussion

Amplitude modulated transcranial alternating current stimulation (AM-tACS) offers a promising new approach to investigate online effects of tACS using physiological recordings. While in theory AM-tACS should not exhibit artifacts within the frequency range of brain signals, the current study demonstrates that non-linear transfer characteristics of stimulation and recording hardware reintroduces such artifacts at the modulation frequency and its lower harmonics. These artifacts are likely too small to modulate brain activity themselves, they can potentially be misinterpreted as stimulation effects on the brain if not considered during concurrent recordings of brain activity during AM-tACS. Especially, in cases where spatial information is missing (i.e. recording from only few EEG sensors), the artifacts in the spectrum might be hard, if not impossible, to be disentangled from stimulation effects. Consequently, these recordings must not be considered artifact-free in the range of the modulation frequency. Rather, the extent of low-frequency artifacts has to be evaluated carefully and taken into account. The setups evaluated for the current study have been build based on a limited set of hardware. Thus, the extent of non-linearity might differ for hardware combinations using other stimulator or recording systems. However, since all electronic components exhibit some degree of non-linearity (Maas, Stephen, 2003), the general process underlying the generation of low-frequency AM-tACS artifacts is potentially applicable to all setups. Only the size of these artifacts can differ depending on the (non-)linearity of the system. The current study provides a framework to measure and estimate a setups transfer characteristics and evaluate the strength of these low-frequency artifacts arising from its non-linearities. Interestingly, the DAC itself exhibited comparatively weak artifacts, while the more complex setups showed stronger artifacts at the modulation frequency and several harmonics. This might indicate that the effect is driven by non-linearities of the stimulator or recording hardware rather than the DAC as suggested by previous authors (Minami and Amano, 2017).

To obtain a model of each setups transfer characteristics, polynomial regression models were fitted to the probe-signal recordings. The degree of the models is a best guess to tradeoff sufficient complexity to capture each setups nonlinearity, and simplicity to retain a straightforward, interpretable model. Unfortunately, traditional approaches for model selection, i.e. based on adjusted *R^2^* or Akaike Information Criterion, that start from a simple intercept or a saturated model, are not applicable to the data at hand, as the non-linearities observed in the setups are very subtle. A simple linear model would already account for a huge proportion of the input-output recordings variance. Adding additional higher degree terms to the model does not sufficiently increase the explained variance to counteract the penalty implemented in most model evaluation metrics. However, as seen in the simulated data only these terms account for the low-frequency artifacts observed in the AM-tACS recordings.

Given that the low-frequency AM-tACS artifacts are several orders of magnitude smaller than the artifact arising during classical tACS (or at the carrier frequency), they are potentially easier to correct/suppress i.e. by applying beamforming (Chander et al., 2016; Witkowski et al., 2016) or temporal signal space separation (Minami and Amano, 2017; Taulu et al., 2005) in the MEG and independent or principal component analysis (ICA/PCA) in the EEG (Helfrich et al., 2014). However, the efficiency of these methods in the context of AM-tACS needs to be systematically investigated in future studies. The optimal solution to overcome the artifacts observed here would be the optimization of stimulation and recording hardware with respect to their linearity. Neither have tES devices currently available been purposefully designed to apply AM-tACS, nor are recording systems for brain activity intended to record AM-signals at intensities as observed during AM-tACS. Devices exhibiting more linear transfer characteristics as i.e. observed for the DAC output in setup 1 would decrease the size of the artifacts compared to the signal of interest such that its influence eventually becomes negligible. Until such devices are available, careful analysis procedures have to be carried out, to ensure trustworthy results from concurrent AM-tACS-M/EEG studies. With the current study an analysis framework is provided that enables researchers to check their AM-tACS setups for non-linearities and spurious low-frequency artifacts and may help to disentangle actual effects of the stimulation on the brain from artifacts introduced by the stimulation.

## 6 Author contributions

FK, EN, FF and CSH, conceived the study. FK collected and analyzed the data. All authors wrote the manuscript.

## 7 Acknowledgements

This research was supported by the Neuroimaging Unit of the Carl von Ossietzky University Oldenburg funded by grants from the German Research Foundation (3T MRI INST 184/152-1 FUGG and MEG INST 184/148-1 FUGG). Christoph S. Herrmann was supported by a grant of the German Research Foundation (DFG, SPP, 1665 3353/8-2). This work was supported in part by the National Institute of Mental Health of the National Institutes of Health under award numbers R01MH111889 and R01MH101547 (PI: Flavio Frohlich). The content is solely the responsibility of the authors and does not necessarily represent the official views of the National Institutes of Health.

## Highlights

- Amplitude modulated tACS generates spurious artifacts at its modulation frequency
- The input-output transfer functions of different AM-tACS setups was estimated
- Hardwares non-linear transfer characteristics account for these spurious artifacts
- An analysis approach to characterize non-linearities of tACS setups is provided.

## Supplementary Materials: Non-linear transfer characteristics of stimulation and recording hardware account for spurious low frequency artifacts during amplitude modulated transcranial alternating current stimulation (AM-tACS)

**Supplementary Figure S1:**
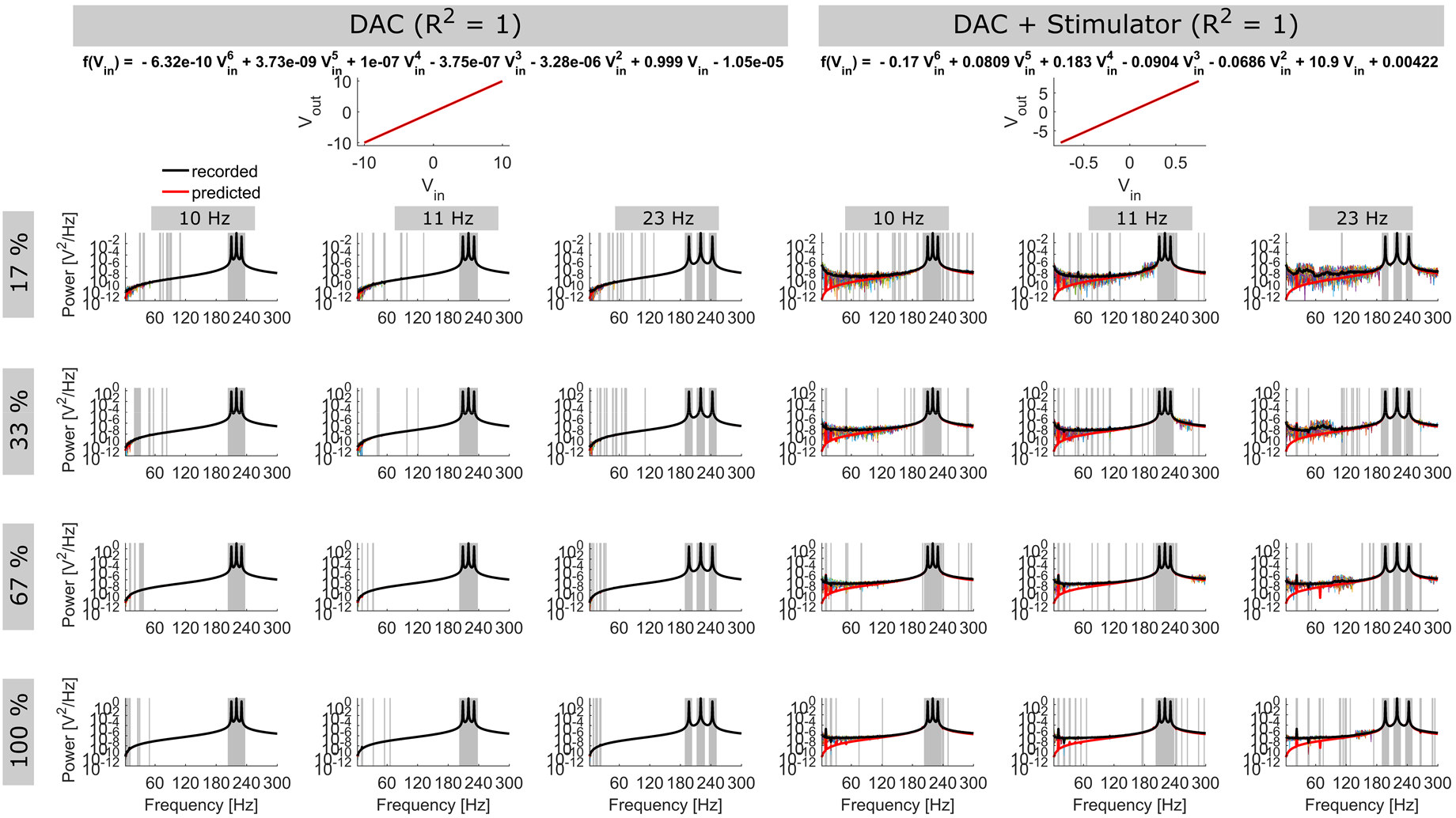
Supplementary Figure S1: Full range version of Figure 2. TFs (top) show recorded probe stimulus amplitudes in relation to their input amplitudes *(V_out_/V_in_*; black dots), as well as the course of the TF model (red line). The corresponding function is displayed in the title. Spectra show average power at each frequency in the different AM-recordings (black line). Thin colored lines show power spectra for each of the 60 repetitions. Red line shows the spectrum predicted by evaluating the digital AM-signal by the estimated TF of the setup. Grey areas indicate frequencies significantly differing in power compared to the two neighboring frequencies (p < .05, bonferroni corrected). Please note the different scaling of the power spectra.

**Supplementary Figure S2:**
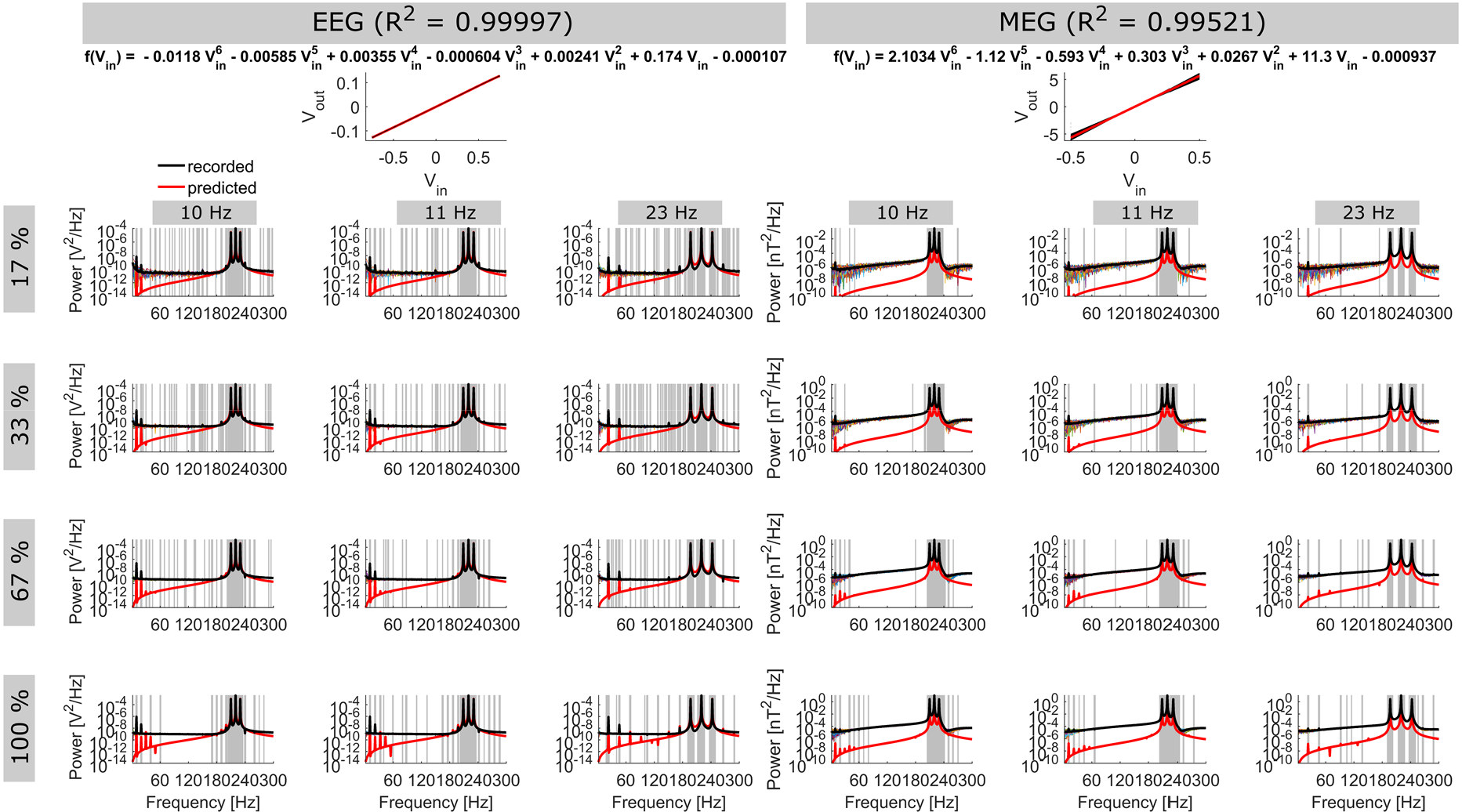
Supplementary Figure S2: Full range version of Figure 3. TFs **(top)** show recorded probe stimulus amplitudes in relation to their input amplitudes *(V_out_/V_in_* black dots), as well as the course of the TF model (red line). The corresponding function is displayed in the title. Spectra show average power at each frequency in the different AM-recordings (black line). Thin colored lines show power spectra for each of the 60 repetitions. Red line shows the spectrum predicted by evaluating the digital AM-signal by the estimated TF of the setup. Grey areas indicate frequencies significantly differing in power compared to the two neighboring frequencies (p < .05, bonferroni corrected). Please note the different scaling of the power spectra.

**Supplementary Figure S3:**
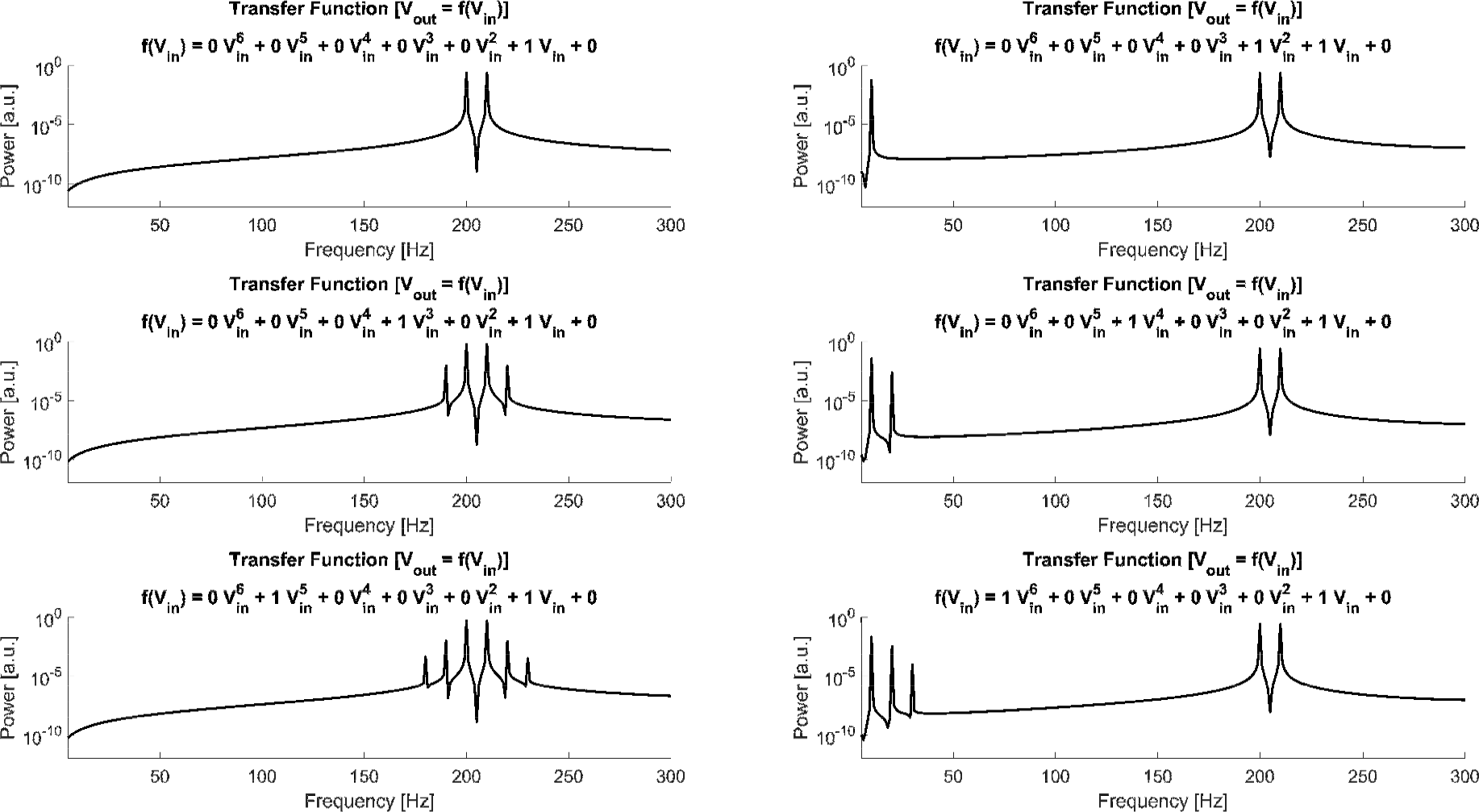
Supplementary Figure S3: Simulation of artifacts resulting from temporal interference (TI). Frequency spectra showing the effect of non-linear TF terms on amplitude modulated signals created by TI. Similar to the am-signals, the TI signals contain no low-frequency artifact when a solely linear TF is applied **(top left).** Adding non-linear terms to the TF model results in additional side-bands around the frequencies of the two applied sine wave signals for odd-valued exponents **(left column)** and in low-frequency artifacts at *Δf* (corresponding to the modulation frequency of the am-signal generated by the TI signals) and its harmonics for even valued exponents of the TF model **(right column).**

## Footnotes

1 In contrast to the frequency-domain definition of TFs commonly used in linear-system analysis, here TF refers to the input-output amplitude relation of a probe signal.

